# Periplasmic carbonic anhydrase CAH1 contributes to high inorganic carbon affinity in *Chlamydomonas reinhardtii*

**DOI:** 10.1101/2024.03.04.583368

**Authors:** Daisuke Shimamura, Tomoaki Ikeuchi, Yoshinori Tsuji, Hideya Fukuzawa, Takashi Yamano

**Author notes:** Author for communications (T.Y.). Senior author (Hideya Fukuzawa).

## Abstract

Carbonic anhydrase (CA), an enzyme conserved across species, is pivotal in the interconversion of inorganic carbon (Ci; CO_2_ and HCO_3_^−^). Compared to the well-studied intracellular CA, the specific role of extracellular CA in photosynthetic organisms is still not well understood. In the green alga *Chlamydomonas reinhardtii*, CAH1, located at the periplasmic space, is strongly induced under CO_2_-limiting conditions by the Myb transcription factor LCR1. While it has been observed that the *lcr1* mutant shows decreased Ci-affinity, the detailed mechanisms behind this phenomenon are yet to be elucidated. In this study, we aimed to unravel the LCR1-dependent genes essential for maintaining high Ci-affinity. To achieve this, we identified a total of 12 LCR1-dependent inducible genes under CO_2_-limiting conditions, focusing specifically on the most prominent ones - *CAH1*, *LCI1*, *LCI6*, and *Cre10.g426800*. We then created mutants of these genes using the CRISPR-Cas9 system, all from the same parental strain, and compared their Ci-affinity. Contrary to earlier findings (Van and Spalding, 1999) that reported no reduction in Ci-affinity in the *cah1* mutant, our newly created *cah1*-1 mutant exhibited a significant decrease in Ci-affinity under high HCO_3_^−^/CO_2_-ratio conditions. Additionally, when we treated wild-type cells with a CA inhibitor with low membrane permeability, a similar reduction in Ci-affinity was observed. Moreover, the addition of exogenous CA to the *cah1* mutant restored the decreased Ci-affinity. These results, highlighting the crucial function of the periplasmic CAH1 in maintaining high Ci-affinity in *Chlamydomonas* cells, provide new insights into the functions of periplasmic CA in algal carbon assimilation.

**One-sentence summary:** CAH1, a periplasmic carbonic anhydrase in *Chlamydomonas reinhardtii*, plays a crucial role in maintaining a high affinity for inorganic carbon, particularly under CO_2_-limiting conditions.

## Introduction

Carbonic anhydrase (CA; EC 4.2.1.1) is a metalloenzyme that catalyzes the interconversion between CO_2_ and HCO_3_^−^. CA is among the enzymes that display the highest turnover rates (Chegwidden and Carter, 2000), thereby fulfilling biological demands in diverse physiological processes such as pH homeostasis, inorganic carbon (Ci: CO_2_ and HCO_3_^−^) transport, and Ci assimilation. CA is classified into nine subclasses (α- to ι-type) based on the primary structure (Aspatwar et al., 2022).

In land plants, CA is hypothesized to play a crucial role in carbon assimilation, although its function remains controversial. Historically, it has been posited that in C3 plants, abundant CA in the chloroplast stroma aids CO_2_ fixation by facilitating its diffusion (Jacobson et al., 1975). However, this understanding has been challenged by recent studies. For instance, Hines et al. (2021) found that the complete loss of stromal CA does not significantly alter carbon assimilation compared to wild-type (WT) plants. In contrast, CAs’ role in aquatic organisms, such as microalgae and cyanobacteria, is more clearly defined within the operation of the CO_2_-concentrating mechanism (CCM) (Fukuzawa et al., 1992, Badger, 2003). Notably, Rubisco, a key enzyme in photosynthetic CO_2_ fixation, exhibits a lower affinity for CO_2_ in microalgae and cyanobacteria than in its terrestrial counterparts (Jordan and Ogren, 1981). To compensate for Rubisco’s lower affinity for CO_2_, particularly in microalgae, the CCM actively transports HCO_3_^−^ into the chloroplast stroma through membrane transporters and channels. Once in the proximity of Rubisco, this HCO_3_^−^ is converted into CO_2_ by intracellular CA, effectively concentrating CO_2_ where it is needed for photosynthesis (Raven et al., 2011).

In *Chlamydomonas reinhardtii*, a freshwater green alga, CAs are crucial for driving CCM, and the compartmentalized CAs are integral to supplying CO_2_ specifically to the pyrenoid, where Rubisco is densely packed in the chloroplast (Moroney et al., 2011). Among them, CAH3, an α-type CA, plays a unique role. It is localized in the lumen of the pyrenoid-invading thylakoid membrane, also known as the pyrenoid tubule. Here, CAH3 facilitates the conversion of HCO_3_^−^ to CO_2_, a process enhanced by the lumen’s acidic pH. CAH3-deficient mutant exhibits decreased Ci affinity with higher accumulation of internal Ci relative to WT cells, highlighting its role in the generation of CO_2_ from the stromal Ci pool (Funke et al., 1997; Karlsson et al., 1998). Additionally, the Low-CO_2_ Inducible Protein B (LCIB), a θ-type CA positioned around the pyrenoid, serves to reconvert CO_2_ leaking from the pyrenoid into HCO_3_^−^, maintaining optimal Ci concentration for photosynthesis (Wang and Spalding, 2006; Yamano et al., 2010; Kasili et al., 2023). In addition to LCIB and CAH3, *Chlamydomonas* has α-type CAs (CAH1–2), β-type CAs (CAH4–9), and γ-type CAs (CAG1–3), but their roles in CCM remain unresolved (Moroney et al., 2011).

CAH1, an α-type CA localized at the periplasmic space, is the first CA to be identified, but its importance in the CCM remains controversial. CAH1 is induced upon CO_2_-limitation, and its induction is dependent on a Myb transcription factor LCR1, whose expression is regulated by CCM1/CIA5, a master regulator of CCM (Fukuzawa et al., 1990, 2001; Xiang et al., 2001; Yoshioka et al., 2004). In addition to the abundant accumulation of CAH1 under CO_2_-limiting conditions, inhibition of periplasmic CA by weakly permeable CA inhibitors, such as acetazolamide (AZA), decreased Ci affinity (Moroney et al., 1985). These data led to the establishment of a well-known model in which periplasmic CA facilitates diffusive CO_2_ entry by maintaining a CO_2_ gradient across the plasma membrane through the rapid equilibration of CO_2_ with bulk HCO_3_^−^ at the cell surface. Because periplasmic CA activity was detected in diverse algae (Nimer et al., 1999; Elzenga et al., 2000; Tsuji et al., 2017, 2021), and its inhibition by AZA caused the decline of Ci-affinity, periplasmic CA-mediated CO_2_ uptake has been a widespread hypothesis. Conversely, it has also been suggested that the effects of AZA on photosynthetic kinetics may be due to the inhibition of intracellular CAs rather than extracellular ones (Williams and Turpin, 1987). Moreover, a *Chlamydomonas* mutant lacking CAH1 showed no difference in growth or Ci-affinity difference under low-CO_2_ conditions (Van and Spalding, 1999), challenging the hypothesis that periplasmic CA facilitates CO_2_ acquisition from bulk HCO_3_^−^. Another hypothesis based on the mathematical modeling is that periplasmic CA recaptures leaked CO_2_ through hydration reaction (Fridlyand, 1997). Thus, while massive effort has been spent to elucidate the function of periplasmic CA, conclusive evidence to support either hypothesis has not been presented yet.

We previously demonstrated through macroarray analysis, which is limited to a specific number of genes, that the *lcr1* mutant was unable to induce at least three LC-inducible genes, namely *CAH1*, *LCI1*, and *LCI6* (Yoshioka et al., 2004). Notably, LCI1, localized at the plasma membrane, is hypothesized to function as a CO_2_ channel due to its structural characteristics (Kono et al., 2020; Ohnishi et al., 2010;). In addition, LCI1 interacts with high-light activated protein 3 (HLA3), an HCO_3_^−^ transporter on the plasma membrane (Yamano et al., 2015; Mackinder et al., 2017). Although the *lcr1* mutant shows a decrease of Ci-affinity under the condition, the major contributor to this phenotype has not been determined yet. Among the three candidates, independent disruption of *CAH1* and *LCI1* do not show decreased Ci-affinities (Van and Spalding, 1999; Kono and Spalding, 2020), suggesting that cooperative functions of these three components or contribution of other unidentified factors for high-affinity photosynthesis for Ci. In this study, to gain further insight into *LCR1*-dependent CCM factors, we generated *lcr1* mutant and identified novel LCR1-dependent genes by RNA-seq analysis. Furthermore, by generating mutant strains of *LCR1*-dependent genes using the CRISPR-Cas9 method, we found that loss of CAH1 causes a decrease in Ci affinity.

## Results

### Identification of LCR1-dependent inducible genes under CO_2_-limiting conditions

In our previous study (Yoshioka et al., 2004), we utilized the *lcr1* insertion mutant derived from the parental strain Q304P3, where *CAH1*-promoter activity was monitored by arylsulfatase (Ars) enzyme activity (Kucho et al., 1999). However, due to the absence of a cell wall, this strain was unsuitable for physiological analysis. To address this, we generated a new *lcr1* mutant, named *lcr1*-1 in this study, derived from the WT strain C9. To create the *lcr1*-1 mutant, we inserted the *AphVII* gene cassette, conferring hygromycin resistance, into the 1^st^ exon of *LCR1* using the CRISPR-Cas9 system (Supplemental Fig. S1A-B). In the *lcr1*-1, the accumulation levels of CAH1 and LCI1 were reduced but recovered in the complemented strain (C-LCR1), where the *LCR1* gene fragment was reintroduced into *lcr1*-1 (Fig. 1). On the other hand, the accumulation levels of HLA3, LCIA, LCIB, LCIC, CAH3, and CCM1 did not change significantly among the strains, consistent with previous findings that LCR1 specifically regulates *CAH1* and *LCI1* among CCM-related genes (Yoshioka et al., 2004).

**Figure 1.**
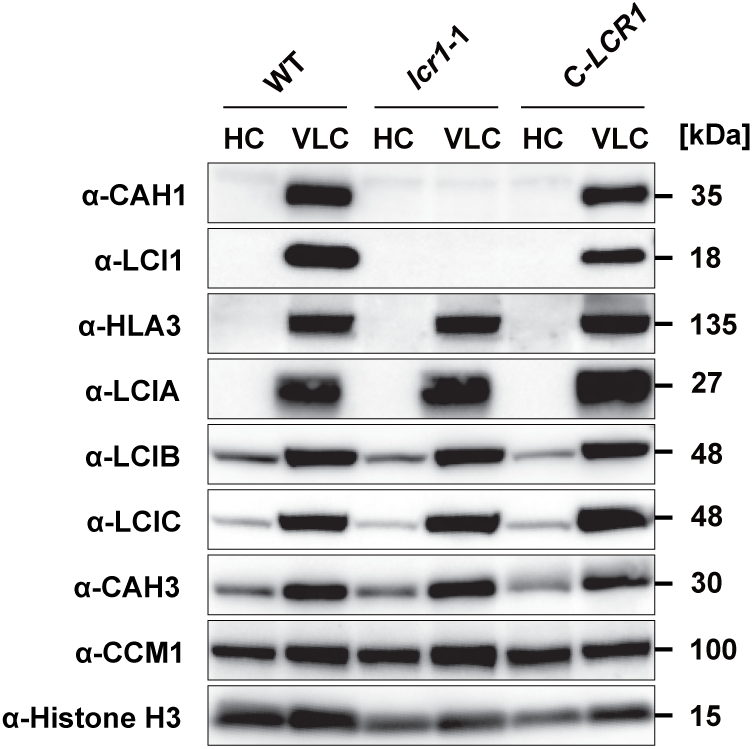
Accumulation of CCM-related proteins in the *lcr1* mutant. Cells were first grown under 5% (v/v) CO_2_ condition for 24 h and shifted to 5% (v/v) CO_2_ (HC) or 0.04% (v/v) (VLC) CO_2_ conditions for 12 h. Histone H3 was used as a loading control.

Given the limitations of macroarray analysis in previous studies for quantifying all gene expression levels (Yoshioka et al., 2004), we explored whether LCR1 influences genes beyond *CAH1*, *LCI1*, and *LCI6* under CO_2_-limiting conditions using RNA-seq analysis. We cultured WT, *lcr1*-1, and C-*LCR1* cells under 5% CO_2_ or 0.04% CO_2_ aerated conditions and quantified their transcriptome profiles. In WT cells, the expression levels of 1,647 genes were significantly increased at either 0.3 or 2 h after switching to LC conditions compared to HC conditions (Supplemental Dataset 1). Among them, under LC conditions, the expression levels of 12 genes, including *LCR1*, *CAH1*, *LCI1*, and *LCI6*, were significantly decreased in *lcr1*-1 and recovered in C-*LCR1* (Table 1). Additionally, the expression level of *Cre10.g426800*, encoding a protein of unknown function with a transferase domain, was decreased more than four-fold by *LCR1* mutation both 0.3 h and 2 h after LC induction (Table 1). This led us to focus our subsequent analysis on *CAH1*, *LCI1*, *LCI6,* and *Cre10.g426800*.

**Table 1.**
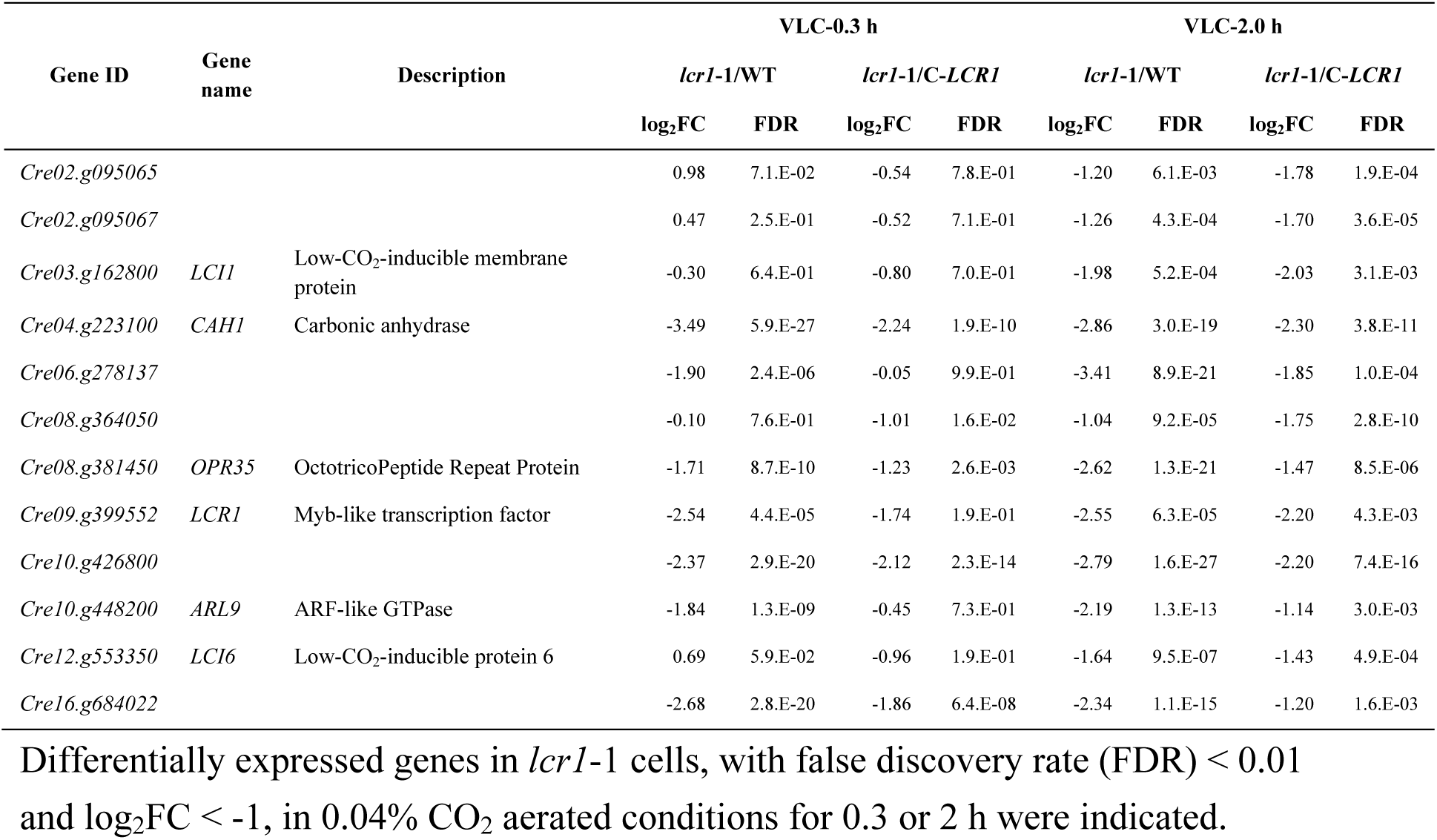
Genes downregulated in the *lcr1* mutant under CO_2_-limiting conditions.

### Impact of CAH1 and LCI1 mutations on Ci-affinity in *Chlamydomonas* cells

To clarify the contribution of LCR1-dependent genes to CCM, we employed the CRISPR-Cas9 method to generate mutants of *CAH1*, *LCI1*, *LCI6*, and *Cre10.g426800*. First, we created insertional mutants of *LCI6* and *Cre10.g426800* and measured their photosynthetic O_2_ evolution rates. Three strains with an insertional mutation in the first exon of *LCI6* were isolated. Additionally, two strains of *Cre10.g426800* were isolated: one with a mutation in the 1^st^ exon and the other in the 2^nd^ exon (Supplemental Fig. S2A-B). To evaluate Ci-affinity in these mutants, we measured their O_2_-evolving activity. The selected pH conditions of 6.2, 7.0, and 7.8 represent a range that encompasses typical environmental variations, allowing us to assess the mutants’ responses under diverse but relevant scenarios. In these mutants, the K_0.5_ (Ci) values, the Ci concentrations required for half-maximal O_2_-evolving rate, showed no significant increase compared to the WT (Supplemental Table S1), suggesting that these two genes were not important for maintaining high Ci-affinity.

Next, we isolated mutants of *CAH1* and *LCI1*, designated *cah1*-1 and *lci1*-1, respectively, and also produced a double mutant (*lci1*/*cah1*-1) by disrupting the *CAH1* in the *lci1*-1 background (Supplemental Fig. S3A-B). The K_0.5_ (Ci) values of the mutants were similar to those of the WT at pH 6.2 and 7.0, with significant differences emerging only at pH 7.8. At this pH, the K_0.5_ (Ci) value of *lcr1*-1 was notably higher than WT, more than four-fold, indicating a reduced Ci-affinity (Fig. 2A and Supplemental Table S1). Interestingly, the *cah1*-1 mutant showed similar K_0.5_ (Ci) values to *lcr1*-1, while the K_0.5_ (Ci) value of *lci1*-1 did not significantly differ from WT. Additionally, the *lci1*/*cah1*-1 double mutant exhibited K_0.5_ (Ci) values comparable to *cah1*-1 and *lcr1*-1, highlighting a critical role for CAH1 in maintaining Ci affinity under a high HCO_3_^−^/CO_2_ ratio. Unexpectedly, we observed a reduction in HLA3 accumulation in *cah1*-1, *lci1*-1, and *lci1*/*cah1*-1 (Fig. 2B). This reduction in HLA3 accumulation complicates our understanding of the roles of CAH1 and HLA3 in maintaining Ci-affinity. It raises the question of whether the observed decrease in Ci-affinity in *cah1*-1 is solely due to CAH1 loss or if it might also involve a synergistic effect resulting from the simultaneous reduction of both CAH1 and HLA3.

**Figure 2.**
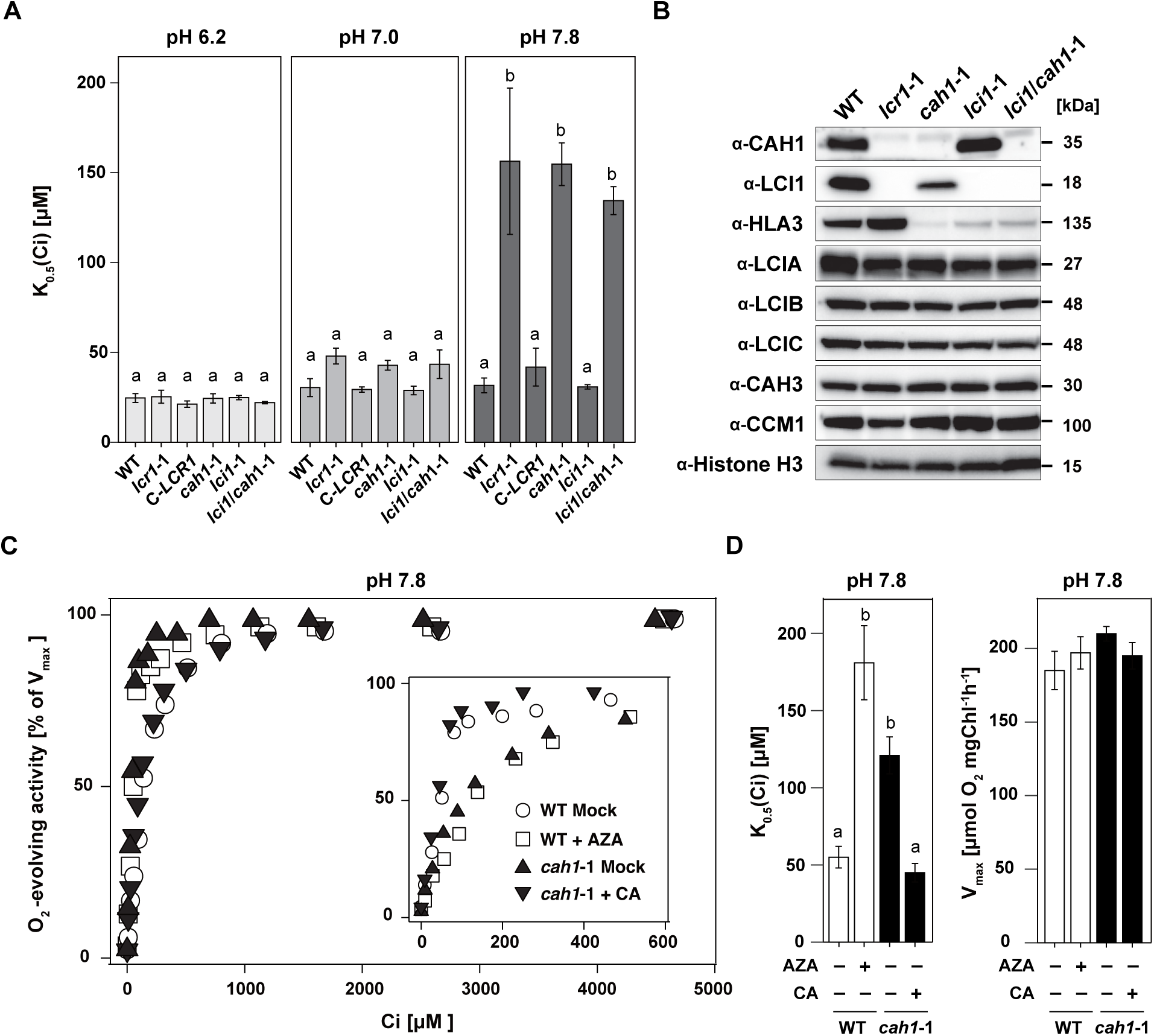
Physiological characteristics of *lcr1*, *cah1*, *lci1* and *lci1*/*cah1* mutants. **A**, K_0.5_ (Ci) values of *cah1*-1, *lci1*-1 and *cah1*/*lci1*-1 cells grown in 0.04% (v/v) CO_2_ conditions for 12 h at pH 6.2, 7.0 and 7.8. Data from all experiments show mean values ± SD from three biological replicates. Statistical analysis was conducted using the Tukey–Kramer multiple comparison test, with different letters indicating significant differences (P < 0.05). **B**, Accumulation of CCM-related proteins in WT, *lcr1*-1, *cah1*-1, *lci1*-1 and *cah1*/*lci1*-1 mutants. Cells were first grown under 5% (v/v) CO_2_ condition for 24 h and shifted to 5% (v/v) CO_2_ or 0.04% (v/v) CO_2_ conditions for 12 h. Histone H3 was used as a loading control. **C**, Typical responses of net O_2_-evolving activities of WT (open circles), WT treated with Acetazolamide (open squares), *cah1*-1 (closed triangle), and *cah1*-1 supplemented with bovine CA (closed inverted triangle) against Ci concentrations at pH 7.8 for the ranges of 0–5000 μM Ci and 0–600 μM Ci (inset). Before measurements, cells were grown in the liquid culture aerated with 0.04% CO_2_ for 12 h. **D**, V_max_ and K_0.5_ (Ci) values of WT treated with AZA and *cah1*-1 supplemented with bovine CA. AZA adjusted to a concentration of 5 mM and dissolved in DMSO, was added to the measuring buffer at a 1% [v/v]. Bovine CA was added into the buffer, achieving a concentration of 2.0 μg/mL. For comparison, 1% DMSO was introduced to samples without AZA. The Tukey–Kramer multiple comparison test was utilized for statistical analysis, with differing letters indicating statistically significant variations (P < 0.05).

### Acetazolamide’s influence on CAH1-mediated Ci-affinity

To further elucidate CAH1’s specific contribution to Ci-affinity and separate its effects from those of HLA3, we investigated the response of cells treated with acetazolamide (AZA), a CA inhibitor with low membrane permeability, at pH 7.8 (Fig. 2C). The addition of AZA to WT cells resulted in an increased K_0.5_ (Ci) value, aligning with the levels observed in *cah1*-1 mutants (Fig. 2D and Supplemental Table S2). Conversely, when *cah1*-1 mutants were supplemented with bovine CA, their K_0.5_ (Ci) value was reduced to WT levels. On the other hand, V_max_ values which indicates the maximum O_2_-evolving activities, did not change among all samples. This demonstrates that the alteration in Ci-affinity observed in *cah1*-1 is primarily attributed to the loss of CAH1 activity rather than HLA3. These results affirm the critical function of periplasmic CAH1 in CCM by maintaining high-Ci affinity under CO_2_-limiting conditions.

### Effect of CAH1 mutation on growth rate

To further examine the impact of CAH1 deficiency, we evaluated the growth rates of WT, *lcr1*-1, and *cah1*-1 cells. Despite the significant role of CAH1 in maintaining Ci-affinity, no differences in growth rate among these strains were observed under both high-CO_2_ (HC) and very low-CO_2_ (VLC) conditions, as evidenced by spot tests on agar plates at pH 7.8 (Fig. 3A). Additionally, the doubling times for these strains in a liquid medium were comparable (Fig. 3B). These findings suggest that, while CAH1 is crucial for maintaining Ci-affinity, its absence does not impede the overall growth rate under CO_2_-limiting conditions.

**Figure 3.**
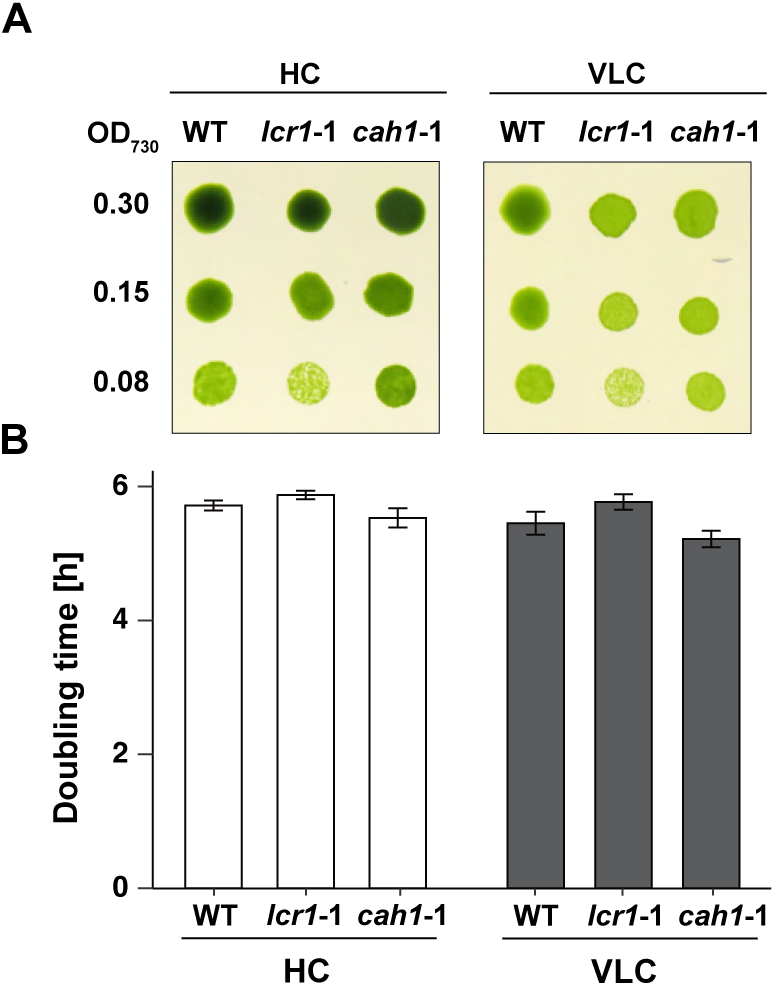
The growth of *lcr1* and *cah1* mutants. **A**, Spot test of WT, *lcr1*-1 and *cah1*-1. Cells were diluted to the indicated optical density (OD_700_ = 0.30, 0.15, or 0.08). Subsequently, 3 μL of the cell suspensions were spotted on agar plates with pH 7.8. The plates were incubated for 4 days under 5% [v/v] CO_2_ (HC) or 0.01% [v/v] CO_2_ (VLC) conditions with continuous light at 120 μmol photons m^-2^ s^-1^ **B**, Doubling time of WT, *lcr1*-1 and *cah1*-1 cells were calculated. Each cell was cultured in a 5% CO_2_ (HC) or 0.04 CO_2_ (VLC) aeration.

## Discussion

In this study, we assessed the Ci-affinity of LCR1-dependent gene mutants created using the CRISPR-Cas9 system across various pH conditions, ranging from acidic to alkaline, to understand their behavior under different environmental scenarios. Notably, at pH 7.8, a condition representative of high HCO_3_^−^/CO_2_ ratios, the *cah1*-1 mutant exhibited a significant decrease in Ci-affinity, highlighting the pivotal role of CAH1 in *Chlamydomonas* cells.

### The function of LCR1 in various environmental stresses

LCR1 is instrumental in the activation of the CCM under CO_2_-limiting conditions, notably regulating the expression of *CAH1* and *LCI1* (Yoshioka et al., 2004). Conversely, under high-light conditions, LCR1 is critical for the expression of *LHCSR3*, essential for photoprotection (Arend et al., 2023). However, our study revealed that LCR1 did not regulate *LHCSR3* expression under CO_2_-limiting conditions (Table 1), demonstrating that the genes controlled by LCR1 vary with environmental context. This highlights the versatile regulatory functions of transcription factors like LCR1, which adapt to different environmental stresses. Such findings underscore the importance of phenotypic analysis under various environmental conditions. Further insights into the diverse functions of transcription factors are expected from the recent large-scale systematic analysis (Fauser et al., 2022), which examines mutant phenotypes under various environmental growth conditions and chemical treatments.

### CAH1 facilitates indirect HCO_3_^−^ utilization under alkaline conditions

We demonstrated that CAH1 is crucial for maintaining high Ci-affinity in *Chlamydomonas* WT cells under alkaline conditions (pH 7.8), supporting the contribution of CAH1 to indirect utilization of abundant HCO_3_^−^ (Fig. 4). This aligns with previous reports indicating enhanced transcription, protein accumulation, and CA activity of CAH1 at higher pH levels (Fett and Coleman, 1994). An earlier study did not reveal significant differences in Ci-affinity between WT and *cah1* mutants (Van and Spalding, 1999), possibly due to the measurements performed at neutral pH. In addition, our research utilized a consistent parental strain for *cah1* mutants, ensuring a more accurate evaluation of CAH1’s impact. As demonstrated in our previous studies (Toyokawa et al, 2020; Tsuji et al., 2023), this study reaffirms the importance of using mutants generated from the same parental strain for accurate phenotypic analysis in *Chlamydomonas* reverse genetics.

**Figure 4.**
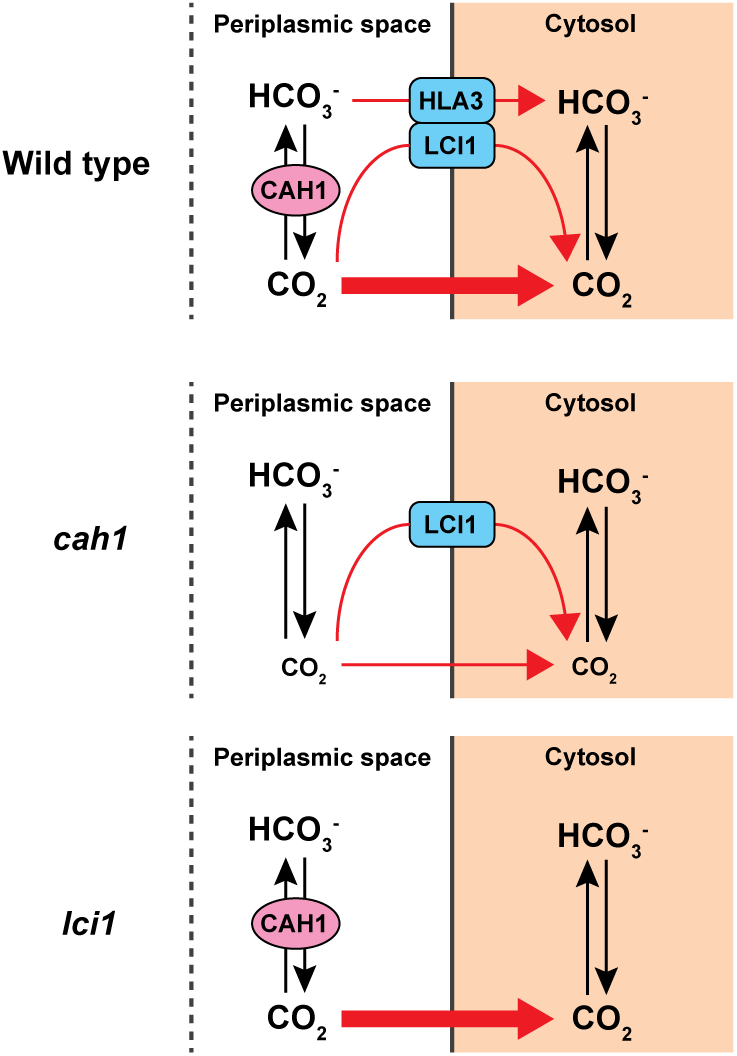
Models for Ci uptake pathway in WT, cah1 and lci1 mutants. Tentative models show how WT, *cah1*, and *lci1* mutants uptake Ci across the plasma membrane under CO_2_-limiting conditions and at pH 7.8. Black arrows indicate the interconversion between CO_2_ and HCO_3_^−^. Red arrows show the Ci uptake pathway from the periplasmic space into the cytosol.

In *Chlamydomonas*, periplasmic CA was identified about four decades ago (Kimpel et al., 1983), and physiological experiments using weakly permeable sulfonamide inhibitors established the well-known model that periplasmic CA facilitates the indirect utilization of bulk HCO_3_^−^ (Moroney et al 1985, Aizawa & Miyachi, 1986). Although the contradictory result in the previous analysis of *cah1* mutant (Van and Spalding, 1999) had raised controversy about the function of periplasmic CA, we demonstrated the importance of CAH1 at alkaline conditions, strengthening the original hypothesis that periplasmic CA supplies CO_2_ from HCO_3_^−^. Regarding catalytic direction (hydration or dehydration), there is an opposing hypothesis based on a mathematical modeling, in which periplasmic CA recaptures CO_2_ leaked from the cell through the hydration (Fridlyand, 1997). However, this hypothesis is unlikely in *Chlamydomonas* because i) analysis using membrane inlet mass spectrometry (MIMS) detected net CO_2_ uptake, but not CO_2_ efflux, by the cell when external CA was inhibited or removed (Sültemeyer et al., 1989, Shiraiwa et al.,1993), and ii) light-dependent alkalization of medium was observed (Shiraiwa et al.,1993). These physiological measurements were performed at pH 8.0, which is similar to the conditions (pH 7.8) where our *cah1* mutant displayed lower Ci-affinity than the parental strain (Fig.2A). Importance of periplasmic CA was also suggested by the pronounced inhibitory effect of weakly permeable sulfonamide inhibitor at alkaline pH (pH 8.0) (Moroney et al.1985). Thus, the long-standing discrepancy between physiological and genetic evidence has been solved, and both approaches provide consistent support for the original model that periplasmic CA enhances indirect HCO_3_^−^ utilization by accelerating dehydration. Besides *Chlamydomonas*, the enhanced CO_2_ uptake by periplasmic CA-mediated dehydration is also supported in some marine diatoms such as *T. pseudonana* and *O. sinensis* by kinetic analysis of CO_2_ uptake using membrane-inlet mass spectrometry (MIMS) and direct measurement of cell surface pH changes, respectively (Hopkinson et al., 2013; Chrachri et al., 2018), suggesting the generality of the classical model in diverse algal groups. Notably, CAH1 in *Chlamydomonas* is α-type while diatoms have δ- and ζ-type in the periplasmic space (Samukawa et al. 2014), suggesting the convergent evolution of the CCM in different lineages as previously discussed (Matsuda et al., 2017).

### Multiple strategies of Ci-uptake in *Chlamydomonas*

In *Chlamydomonas*, Ci uptake across the plasma membrane involves multiple transport strategies. These include direct pathways of CO_2_ through LCI1 and HCO_3_^−^ through HLA3, respectively, along with indirect pathways involving CAH1 (Fig. 4). Despite no significant reduction in Ci affinity in *lci1*-1 mutants (Fig. 2A and Supplemental Table S1), LCI1’s cooperative role with other transporters cannot be ruled out.

Interestingly, the accumulation level of HLA3 was reduced in *cah1*-1 and *lci1*-1 (Fig. 4). The complete loss of CAH1 and LCI1 may have caused this phenotype, as HLA3 accumulation was not altered in *lcr1*-1 (Fig. 1 and 2B). Since HLA3 and LCI1 interact and form a complex on the plasma membrane (Mackinder et al., 2017), it is possible that the formation of the HLA3-LCI1 complex was inhibited in *lci1*-1. Moreover, the changes in the HCO_3_^−^/CO_2_ ratio in periplasmic space or increased CO_2_ leakage due to CAH1 mutation may have also affected the expression of HLA3. This result suggests a regulatory mechanism for HLA3 expression in response to changes in extracellular Ci.

Notably, from our findings that *cah1*-1 mutants displayed a substantially higher K_0.5_ (Ci) value compared to WT under pH 7.8 conditions (Fig. 2A and Supplemental Table S1), emphasizing CAH1’s primary role in Ci uptake into chloroplasts at this pH conditions (Fig. 4). On the other hands, the absence of significant growth rate differences between *lcr1*-1, *cah1*-1, and WT under CO_2_-limiting conditions suggests the presence of compensatory Ci uptake mechanisms (Fig. 3A and B).

### Diversity of periplasmic CA functions in *Chlamydomonas*

Our findings reveal that AZA significantly reduced Ci-affinity in cells, aligning with previous research (Moroney et al., 1985). Besides CAH1, CAH2, and CAH8 are also located in the periplasmic space (Moroney et al., 2011). Although CAH2 shares a similar amino acid sequence with CAH1, its expression is induced under high CO_2_ conditions, differing from CAH1 (Fujiwara et al., 1990). CAH8, a β-type CA with a transmembrane domain, is positioned closer to the plasma membrane than CAH1 under varying CO_2_ conditions (Ynalvez et al., 2008). The comparable Ci-affinity in AZA-treated WT cells and *cah1* mutant underscores CAH1’s greater role in Ci uptake under CO_2_-limiting conditions compared to CAH2 and CAH8. Future studies focusing on the regulation of these periplasmic CAs and their compensatory interactions are essential, potentially informing bioengineering approaches to enhance microalgae photosynthesis.

## Materials and Methods

### *Chlamydomonas* strains and cultural conditions

The wild-type (WT) strain C9, obtained from the IAM Culture Collection at the University of Tokyo, was utilized for physiological and biochemical experiments. Strain C9 is now available from the Microbial Culture Collection at the National Institute for Environmental Studies, Japan, as strain NIES-2235 (alternatively named CC-5098 in the *Chlamydomonas* Resource Center. The cells were pre-cultured in a TAP medium and subsequently resuspended in 50 mL of MOPS-P medium. They were grown under a 5% (v/v) CO_2_ atmosphere with a light intensity set at 120 μmol photons·m^−2^·s^−1^, following the method described by Toyokawa et al. (2020), until they reached the mid-logarithmic phase of growth. For the induction of very low-CO_2_ (VLC) conditions, cells acclimated to high-CO_2_ conditions were centrifuged, resuspended in fresh MOPS-P medium, and then cultured with air bubbling containing 0.04% (v/v) CO_2_ at the same light intensity.

### Measurement of photosynthetic CO_2_-evolving activity

Cells were harvested and resuspended in Ci-depleted 20 mM MES-NaOH (pH 6.2), MOPS-NaOH (pH 7.0), or HEPES-NaOH (pH 7.8) buffers, adjusting the density to 10– 20 μg chlorophyll per mL. The photosynthetic oxygen evolution rate was then measured using a Clark-type oxygen electrode (Hansatech Instruments).

### Generation of mutants by the CRISPR-Cas9 system

For CRISPR-Cas9-mediated genome editing, guide RNAs were designed using the CRISPOR tool (Concordet and Haeussler, 2018), as detailed in Supplemental Figures S1–3. The introduction of the ribonucleoprotein complex and the *AphVII* or *AphVIII* cassette into cells followed the method of Tsuji et al. (2022).

### Immunoblotting analysis

Total protein extraction, SDS-polyacrylamide gel electrophoresis (SDS/PAGE), and immunoblotting analyses were carried out as previously described (Shimamura et al., 2023). Primary antibodies were utilized at the following indicated dilutions: anti-HLA3 at 1:1,250, anti-LCIA at 1:5,000, anti-LCI1 at 1:5,000, anti-LCIB at 1:5,000, anti-CAH1 at 1:2,500, anti-CAH3 at 1:2,000, anti-CCM1 at 1:2,500, and anti-Histone H3 at 1:10,000. A horseradish peroxidase-conjugated goat anti-rabbit IgG antibody from Life Technologies was employed as the secondary antibody at a dilution of 1:10,000 to detect the primary antibodies.

### RNA-Seq analysis

Total RNA was extracted from cells using the RNeasy Plant Mini Kit (QIAGEN), following the manufacturer’s instructions. After RNA purification, the total RNA was analyzed using the Illumina Novaseq 6000 system. The resulting reads were aligned with version 5.6 of the *Chlamydomonas reinhardtii* genome annotation, which was downloaded from https://phytozome-next.jgi.doe.gov/. The alignment, counting of reads, and normalization of read counts were performed according to the methods previously described in Shimamura et al. (2023).

## ACCESSION NUMBERS

The accession numbers of the Phytozome database for *Chlamydomonas* genes *LCR1*, *CAH1*, *LCI1*, and *LCI6* are *Cre09.g399552*, *Cre04.g223100*, *Cre03.g162800*, and *Cre12.g553350*, respectively.

## Data Availability

Data deposition: The RNA-seq raw data in this paper have been deposited in the DNA Data Bank of Japan (DDBJ) Sequence Read Archive (DRA) (accession no. DRA017670).

## ACKNOWLEDGEMENTS

This work was supported by JSPS KAKENHI Grant Number JP16H06279 (PAGS).

## Author contributions

T.Y. and H.F. conceived and designed the study; D.S. performed most of the experiments; T.I. contributed to mutant isolation; D.S., Y.T., and T.Y. wrote the article, and all authors approved it. T.Y. agreed to serve as the author responsible for contact and ensure communication.

## Author for Contact

The author responsible for distribution of materials integral to the findings presented in this article in accordance with the policy described in the Instructions for Authors (https://academic.oup.com/plphys/pages/general-instructions) is: Takashi Yamano.

## FUNDING

This work was supported by the Japan Society for the Promotion of Science (Grants Numbers JP20H03073 and JP21K19145 to T.Y.), JST SPRING JPMJSP2110 (to D.S.), GteX Program Japan Grant Number JPMJGX23B0 (to T.Y.), and the Asahi Glass Foundation (to T.Y.)

## SUPPLEMENTAL DATA

The following supplemental materials are available.

**Supplemental Figure S1. The *lcr1* mutant generated by the CRISPR-Cas9 system.**

A, The position of guide RNA (gRNA) and protospacer adjacent motif (PAM) sequence in the first exon of *LCR1* gene. gRNA sequences are underlined and PAM sequences are in red letters. The solid rectangles indicate the exons of *LCR1* gene. Arrows indicate primers utilized for PCR screening. B, Genomic PCR to confirm insertion of the *AphVII* cassette in *LCR1* gene.

**Supplemental Figure S2. The mutants of *lci6* and *Cre10.g426800* generated by the CRISPR-Cas9 system.**

A, The positions of guide RNA (gRNA) and protospacer adjacent motif (PAM) sequence in *LCI6* and *Cre10.g426800* gene. gRNA sequences are underlined and PAM sequences are in red letters. The solid rectangles indicate the exons of *LCR1* gene. Arrows indicate primers utilized for PCR screening. B, Genomic PCR to confirm insertion of the *AphVII* cassette in *LCI6* and *Cre10.g426800* gene.

**Supplemental Figure S3. The mutants of *cah1* and *lci1* generated by the CRISPR-Cas9 system.**

A, The positions of guide RNA (gRNA) and protospacer adjacent motif (PAM) sequence in *CAH1* and *LCI1* gene. gRNA sequences are underlined and PAM sequences are in red letters. The solid rectangles indicate the exons of *CAH1* and *LCI1* gene. Arrows indicate primers utilized for PCR screening. B, Genomic PCR to confirm insertion of the *AphVIII* or *AphVII* cassette in *CAH1* and *LCI1* gene. An asterisk indicates a non-specific band.

**Supplemental Table S1. Photosynthetic parameters of WT and transformant cells.**

Cells grown in 5% CO_2_ were shifted to 0.04% CO_2_ for 24 h at 120 μmol photons m^-2^ s^-1^. The data are shown ± standard deviation, which was obtained from three independent experiments. V_max_, maximum O_2_-evolving activity; K_0.5_ (Ci), Ci concentration required for half of V_max_.

**Supplemental Table S2. Effect of AZA and bovine CA on photosynthetic parameters of WT and *cah1*-1 cells.**

Cells grown in 5% CO_2_ were shifted to 0.04% CO_2_ for 24 h at 120 μmol photons m^-2^ s^-1^. The data are shown ± standard deviation, which was obtained from three independent experiments. V_max_, maximum O_2_-evolving activity; K_0.5_ (Ci), Ci concentration required for half of V_max_. In experiment with AZA or bovine CA, 1% (v/v) DMSO was used as a mock.

## REFERENCES

Aizawa K, Miyachi S (1986) Carbonic anhydrase and CO_2_ concentrating mechanisms in microalgae and cyanobacteria. FEMS Microbiol. Rev. 2: 215–233

Arend M, Yuan Y, Ruiz-Sola MÁ, Omranian N, Nikoloski Z, Petroutsos D (2023) Widening the landscape of transcriptional regulation of green algal photoprotection. Nat. Commun. 14: 2687

Aspatwar A, Tolvanen MEE, Barker H, Syrjänen L, Valanne S, Purmonen S, Waheed A, Sly WS, Parkkila S (2022) Carbonic anhydrases in metazoan model organisms: molecules, mechanisms, and physiology. Physiol. Rev. 102: 1327–1383

Badger M (2003) The roles of carbonic anhydrases in photosynthetic CO_2_ concentrating mechanisms. Photosynth. Res. 77: 83–94

Chegwidden WR, Carter ND (2000) Introduction to the carbonic anhydrases. *In* WR Chegwidden, ND Carter, YH Edwards, eds, The Carbonic Anhydrases: New Horizons. Birkhäuser, Basel, pp 13–28

Chrachri A, Hopkinson BM, Flynn K, Brownlee C, Wheeler GL (2018) Dynamic changes in carbonate chemistry in the microenvironment around single marine phytoplankton cells. Nat. Commun. 9: 74

Concordet J-P, Haeussler M (2018) CRISPOR: intuitive guide selection for CRISPR/Cas9 genome editing experiments and screens. Nucleic Acids Res. 46: W242–W245

Elzenga JTM, Prins HBA, Stefels J (2000) The role of extracellular carbonic anhydrase activity in inorganic carbon utilization of *Phaeocystis globosa* (*Prymnesiophyceae*): A comparison with other marine algae using the isotopic disequilibrium technique. Limnol. Oceanogr. 45: 372–380

Fauser F, Vilarrasa-Blasi J, Onishi M, Ramundo S, Patena W, Millican M, Osaki J, Philp C, Nemeth M, Salomé PA, et al (2022) Systematic characterization of gene function in the photosynthetic alga *Chlamydomonas reinhardtii*. Nat. Genet. 54: 705–714

Fett JP, Coleman JR (1994) Regulation of Periplasmic Carbonic Anhydrase Expression in *Chlamydomonas reinhardtii* by Acetate and pH. Plant Physiol. 106: 103–108

Fridlyand LE (1997) Models of CO_2_ concentrating mechanisms in microalgae taking into account cell and chloroplast structure. Biosystems 44: 41–57

Fujiwara S, Fukuzawa H, Tachiki A, Miyachi S (1990) Structure and differential expression of two genes encoding carbonic anhydrase in *Chlamydomonas reinhardtii*. Proc. Natl. Acad. Sci. U. S. A. 87: 9779–9783

Fukuzawa H, Suzuki E, Komukai Y, Miyachi S (1992) A gene homologous to chloroplast carbonic anhydrase (icfA) is essential to photosynthetic carbon dioxide fixation by *Synechococcus* PCC7942. Proc. Natl. Acad. Sci. U. S. A 89: 4437–4441

Fukuzawa H, Fujiwara S, Yamamoto Y, Dionisio-Sese ML, Miyachi S (1990) cDNA cloning, sequence, and expression of carbonic anhydrase in *Chlamydomonas reinhardtii*: regulation by environmental CO_2_ concentration. Proc. Natl. Acad. Sci. U. S. A. 87: 4383–4387

Fukuzawa H, Miura K, Ishizaki K, Kucho K, Saito T, Kohinata T, Ohyama K (2001) Ccm1, a regulatory gene controlling the induction of a carbon-concentrating mechanism in *Chlamydomonas reinhardtii* by sensing CO_2_ availability. Proc. Natl. Acad. Sci. U. S. A. 98: 5347–5352

Funke RP, Kovar JL, Weeks DP (1997) Intracellular carbonic anhydrase is essential to photosynthesis in *Chlamydomonas reinhardtii* at atmospheric levels of CO_2_. Demonstration via genomic complementation of the high-CO_2_-requiring mutant ca-1. Plant Physiol 114: 237–244

Hines KM, Chaudhari V, Edgeworth KN, Owens TG, Hanson MR (2021) Absence of carbonic anhydrase in chloroplasts affects C3 plant development but not photosynthesis. Proc. Natl. Acad. Sci. U. S. A. 118: e2107425118

Hopkinson BM, Meile C, Shen C (2013) Quantification of Extracellular Carbonic Anhydrase Activity in Two Marine Diatoms and Investigation of Its Role. Plant Physiol. 162: 1142–1152

Jacobson BS, Fong F, Heath RL (1975) Carbonic Anhydrase of Spinach: Studies on Its Location, Inhibition, and Physiological Function. Plant Physiol. 55: 468–474

Jordan DB, Ogren WL (1981) Species variation in the specificity of ribulose biphosphate carboxylase/oxygenase. Nature 291: 513–515

Karlsson J, Clarke AK, Chen ZY, Hugghins SY, Park YI, Husic HD, Moroney JV, Samuelsson G (1998) A novel alpha-type carbonic anhydrase associated with the thylakoid membrane in *Chlamydomonas reinhardtii* is required for growth at ambient CO_2_. EMBO J. 17: 1208–1216

Kasili RW, Rai AK, Moroney JV (2023) LCIB functions as a carbonic anhydrase: evidence from yeast and *Arabidopsis* carbonic anhydrase knockout mutants. Photosynth. Res. 156: 193–204

Kimpel DL, Togasaki RK, Miyachi S (1983) Carbonic Anhydrase in *Chlamydomonas reinhardtii* I. Localization. Plant Cell Physiol. 24: 255–259

Kono A, Chou T-H, Radhakrishnan A, Bolla JR, Sankar K, Shome S, Su C-C, Jernigan RL, Robinson CV, Yu EW, et al (2020) Structure and function of LCI1: a plasma membrane CO_2_ channel in the *Chlamydomonas* CO_2_ concentrating mechanism. Plant J. 102: 1107–1126

Kono A, Spalding MH (2020) LCI1, a *Chlamydomonas reinhardtii* plasma membrane protein, functions in active CO_2_ uptake under low CO_2_. Plant J. 102: 1127–1141

Kucho K i, Ohyama K, Fukuzawa H (1999) CO_2_-responsive transcriptional regulation of CAH1 encoding carbonic anhydrase is mediated by enhancer and silencer regions in *Chlamydomonas reinhardtii*. Plant Physiol. 121: 1329–1338

Mackinder LCM, Chen C, Leib RD, Patena W, Blum SR, Rodman M, Ramundo S, Adams CM, Jonikas MC (2017) A Spatial Interactome Reveals the Protein Organization of the Algal CO_2_-Concentrating Mechanism. Cell 171: 133–147.e14

Matsuda Y, Hopkinson BM, Nakajima K, Dupont CL, Tsuji Y (2017) Mechanisms of carbon dioxide acquisition and CO_2_ sensing in marine diatoms: a gateway to carbon metabolism. *Philos. Trans. R. Soc. Lond., B*, Biol. Sci. 372: 20160403

Moroney JV, Husic HD, Tolbert NE (1985) Effect of Carbonic Anhydrase Inhibitors on Inorganic Carbon Accumulation by *Chlamydomonas reinhardtii*. Plant Physiol. 79: 177–183

Moroney JV, Ma Y, Frey WD, Fusilier KA, Pham TT, Simms TA, DiMario RJ, Yang J, Mukherjee B (2011) The carbonic anhydrase isoforms of *Chlamydomonas reinhardtii*: intracellular location, expression, and physiological roles. Photosynth. Res. 109: 133–149

Nimer NA, Brownlee C, Merrett MJ (1999) Extracellular carbonic anhydrase facilitates carbon dioxide availability for photosynthesis in the marine dinoflagellate prorocentrum micans. Plant Physiol. 120: 105–112

Ohnishi N, Mukherjee B, Tsujikawa T, Yanase M, Nakano H, Moroney JV, Fukuzawa H (2010) Expression of a low CO₂-inducible protein, LCI1, increases inorganic carbon uptake in the green alga *Chlamydomonas reinhardtii*. Plant Cell 22: 3105–3117

Raven JA, Giordano M, Beardall J, Maberly SC (2011) Algal and aquatic plant carbon concentrating mechanisms in relation to environmental change. Photosynth. Res. 109: 281–296

Samukawa M, Shen C, Hopkinson BM, Matsuda Y (2014) Localization of putative carbonic anhydrases in the marine diatom, *Thalassiosira pseudonana*. Photosynth. Res. 121: 235–249

Shimamura D, Yamano T, Niikawa Y, Hu D, Fukuzawa H (2023) A pyrenoid-localized protein SAGA1 is necessary for Ca^2+^-binding protein CAS-dependent expression of nuclear genes encoding inorganic carbon transporters in *Chlamydomonas reinhardtii*. Photosynth. Res. 156: 181–192

Shiraiwa Y, Goyal A, Tolbert NE (1993) Alkalization of the Medium by Unicellular Green Algae during Uptake Dissolved Inorganic Carbon. Plant Cell Physiol. 34: 649–657

Sültemeyer DF, Miller AG, Espie GS, Fock HP, Canvin DT (1989) Active CO_2_ Transport by the Green Alga *Chlamydomonas reinhardtii*. Plant Physiol. 89: 1213–1219

Toyokawa C, Yamano T, Fukuzawa H (2020) Pyrenoid Starch Sheath Is Required for LCIB Localization and the CO_2_-Concentrating Mechanism in Green Algae. Plant Physiol. 182: 1883–1893

Tsuji Y, Kinoshita A, Tsukahara M, Ishikawa T, Shinkawa H, Yamano T, Fukuzawa H (2023) A YAK1-type protein kinase, triacylglycerol accumulation regulator 1, in the green alga *Chlamydomonas reinhardtii* is a potential regulator of cell division and differentiation into gametes during photoautotrophic nitrogen deficiency. J. Gen. Appl. Microbiol. 69: 1–10

Tsuji Y, Kusi-Appiah G, Kozai N, Fukuda Y, Yamano T, Fukuzawa H (2021) Characterization of a CO_2_-Concentrating Mechanism with Low Sodium Dependency in the Centric Diatom *Chaetoceros gracilis*. Mar. Biotechnol. (NY) 23: 456–462

Tsuji Y, Mahardika A, Matsuda Y (2017) Evolutionarily distinct strategies for the acquisition of inorganic carbon from seawater in marine diatoms. J. Exp. Bot. 68: 3949–3958

Van K, Spalding MH (1999) Periplasmic carbonic anhydrase structural gene (*Cah1*) mutant in *Chlamydomonas reinhardtii*. Plant Physiol. 120: 757–764

Wang Y, Spalding MH (2006) An inorganic carbon transport system responsible for acclimation specific to air levels of CO_2_ in *Chlamydomonas reinhardtii*. Proc. Natl. Acad. Sci. U. S. A. 103: 10110–10115

Williams TG, Turpin DH (1987) The Role of External Carbonic Anhydrase in Inorganic Carbon Acquisition by *Chlamydomonas reinhardii* at Alkaline pH. Plant Physiol. 83: 92–96

Xiang Y, Zhang J, Weeks DP (2001) The *Cia5* gene controls formation of the carbon concentrating mechanism in *Chlamydomonas reinhardtii*. Proc. Natl. Acad. Sci. U.S. A. 98: 5341–5346

Yamano T, Sato E, Iguchi H, Fukuda Y, Fukuzawa H (2015) Characterization of cooperative bicarbonate uptake into chloroplast stroma in the green alga *Chlamydomonas reinhardtii*. Proc. Natl. Acad. Sci. U. S. A. 112: 7315–7320

Yamano T, Tsujikawa T, Hatano K, Ozawa S-I, Takahashi Y, Fukuzawa H (2010) Light and low-CO_2_-dependent LCIB-LCIC complex localization in the chloroplast supports the carbon-concentrating mechanism in *Chlamydomonas reinhardtii*. Plant Cell Physiol. 51: 1453–1468

Ynalvez RA, Xiao Y, Ward AS, Cunnusamy K, Moroney JV (2008) Identification and characterization of two closely related beta-carbonic anhydrases from *Chlamydomonas reinhardtii*. Physiol. Plant. 133: 15–26

Yoshioka S, Taniguchi F, Miura K, Inoue T, Yamano T, Fukuzawa H (2004) The novel Myb transcription factor LCR1 regulates the CO_2_-responsive gene *Cah1*, encoding a periplasmic carbonic anhydrase in *Chlamydomonas reinhardtii*. Plant Cell 16: 1466–1477

